# Oncogenic IDH mutations increase heterochromatin-related replication stress without impacting tumor mutation burden

**DOI:** 10.1101/2022.08.04.502834

**Authors:** Juan-Manuel Schvartzman, Grace Forsyth, Henry Walch, Walid Chatila, Anthony Santella, Kamal Menghrajani, Francisco Sánchez-Vega, Richard Koche, Craig B. Thompson

## Abstract

Oncogenic mutations in the metabolic enzyme isocitrate dehydrogenase 1 and 2 (IDH1/2) have been found in a number of liquid and solid tumors. Their pathogenic mechanism of action involves production of 2-hydroxyglutarate (2HG), an oncometabolite that acts in part by inhibiting members of a family of dioxygenases that modulate chromatin dynamics. Recent work has suggested that mutant IDH (mIDH) and 2HG also impact sensitivity to inhibitors of poly-ADP ribose polymerases (PARP) but the molecular basis for this sensitivity is unclear. Unlike PARP inhibitor-sensitive BRCA1/2 tumors which exhibit impaired homologous recombination, IDH-mutant tumors have a silent mutational profile and lack mutational signatures associated with impaired homologous recombination. Instead, 2HG-producing IDH mutations lead to heterochromatin-dependent slowing of DNA replication and increased replication stress, resulting in DNA double strand breaks. This replicative stress manifests as replication fork slowing but the breaks are repaired without a significant increase in the cellular mutation burden. Faithful resolution of replicative stress in IDH-mutant cells is dependent on poly-ADP ribosylation. Treatment with PARP inhibitors restores replication fork speed but results in incomplete repair of DNA breaks. These findings provide evidence of a requirement for PARP in the replication of heterochromatin and further validate PARP as a potential therapeutic target in IDH-mutant tumors.

## Introduction

The finding that metabolic rewiring can impact chromatin accessibility has expanded our understanding of how gene expression is regulated. It is now well accepted that metabolic pathways that lead to histone acetylation and deacetylation, DNA and histone methylation and demethylation as well as other metabolite-derived histone modifications, can dramatically impact whether genes are actively expressed or repressed (Dai et al., 2020). How these metabolic reactions can drive locus-specific chromatin changes is unclear, however, and a prevailing model is that metabolism-dependent chromatin processes take place at compartment-size resolution (~10^5^-10^6^ bps).

Oncogenic point mutations in the enzyme isocitrate dehydrogenase (cytoplasmic IDH1 and mitochondrial IDH2) are found in a wide array of tumors and result in a neomorphic enzyme that can reduce alpha-ketoglutarate (aKG) to 2-hydroxyglutarate (2HG) (Amary et al., 2011; Dang et al., 2009; Lu et al., 2012; Ward et al., 2010). 2HG has been found to inhibit the activity of aKG-dependent dioxygenases. Among these dioxygenases are a family of lysine demethylases (KDM) responsible for hydroxylating and removing methylation marks from lysines within histone tails. Thus, IDH mutations have been associated with hypermethylation of histone residues, leading to impaired chromatin accessibility. Tumors harboring IDH mutations show histological evidence of a differentiation block, and this impairment in chromatin accessibility in IDH-mutant cells has been shown to lead to impaired differentiation in model systems (Schvartzman et al., 2019).

An alternative model to how IDH mutations drive oncogenesis was recently proposed based on the observation that IDH-mutant cells showed increased numbers of DNA double-strand-breaks and were sensitive to inhibitors of poly-ADP-ribose polymerases (PARP) (Sulkowski et al., 2017, 2020, 2020). According to this model, homologous recombination requires demethylation of methylated histone H3 lysine 9 residues. In the presence of 2HG, this demethylation reaction is inhibited and homologous recombination is impaired. This impaired homologous recombination is proposed to explain the synthetic lethality with PARP inhibitors, as previously reported for tumors harboring mutations in DNA-damage response genes such as BRCA1 and BRCA2.

Given that the pattern of tumor types harboring mutations in IDH1 and IDH2 was distinct from tumor types harboring BRCA1/2 mutations, we decided to take an in depth look at the mutational patterns observed in IDH1/2 mutant tumors. Here we show that IDH1/2 mutant tumors do not show the typical genomic signatures of impaired homologous recombination. Rather, IDH1/2 mutant tumors exhibit few mutations. However, we found that the increased heterochromatin present in IDH1/2 mutant cells was associated with a prolonged S phase and slower replication fork progression that was associated with increased replicative stress. This replicative stress results in DNA double strand breaks but the breaks are efficiently repaired in a manner that is dependent on poly-ADP-ribose polymerase activity.

## Results

### Analysis of mutation spectrum of IDH-mutated human tumors

To determine whether IDH-mutated tumors shared mutational signatures with tumors with known defects in homologous recombination, we undertook mutational analysis in two independent cohorts of tumor samples for which sequencing data was readily available. Homologous recombination deficiency leads to large scale insertions, deletions and amplifications, and these can be identified through the large scale transition score (LST), a measure of chromosomal breaks between adjacent regions of at least 10 Mb in size (Popova et al., 2012). When we calculated the LST for tumors in the MSK-IMPACT database and subclassified samples according to mutation subclass (BRCA1/2 mutant, other mutations in DNA damage repair genes, IDH1/2 mutant and not mutated for DDR genes) we found that IDH1/2 mutant tumors had a significantly lower LST than either BRCA1/2 or DDR mutant tumors (Figure 1A). This pattern was also present in an independent dataset from The Cancer Genome Atlas Program (TCGA) (Figure 1B) (Popova et al., 2012).

**Figure 1.**
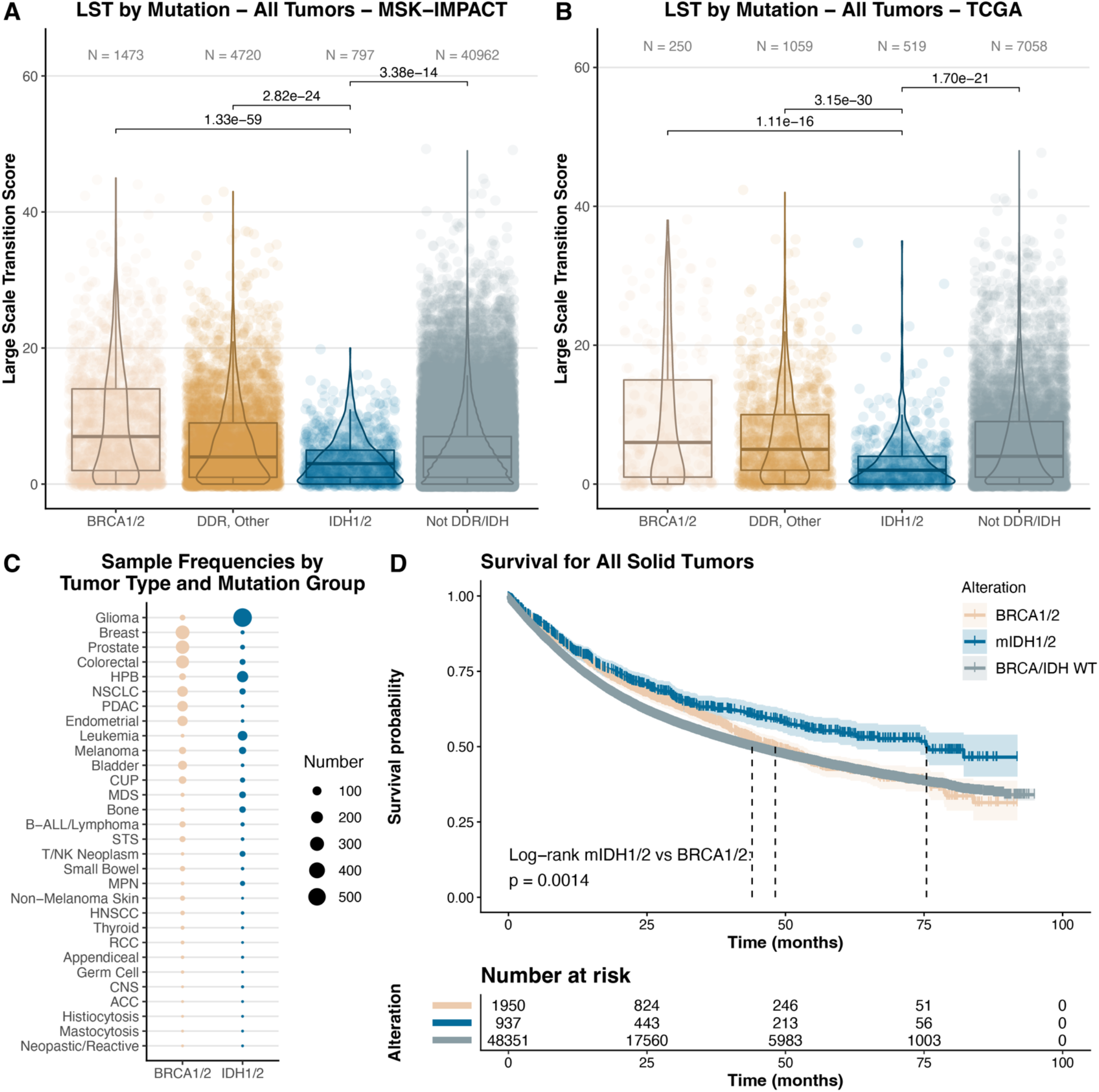
IDH1/2-mutant tumors have a low mutation burden and show improved survival compared to DNA-Damage Response mutations. A. Large Scale Transition (LST) Score from all tumor types in MSK-IMPACT data set classified by mutation status. B. LST from all tumor types in TCGA data set. C. Tumor sample frequencies by tumor type MSK-IMPACT data set by mutation status. D. Survival probability for tumors with the indicated mutation classes. P values for A and B correspond to Dunn’s test after a Kruskal-Wallis with p = 2.5e-87 (A) and p = 1.9e-31 (B). P value for D corresponds to a log-rank test.

IDH1 mutations are frequently observed in lower grade gliomas (Figure 1C) and hence overrepresentation of these tumors could explain the lower LST scores. However, the LST score in IDH1/2-mutant tumors still remained significantly lower than BRCA1/2 or DDR-deficient tumors when gliomas were omitted from the analysis (Supplementary Figure 1A-B). If the analysis was focused on subsets of tumor types seen in all three analyzed groups, IDH1/2-mutated tumors still had significantly lower LST scores. This was true in melanoma, sarcoma, non-small cell lung cancer, hepatobiliary and prostate cancer (Supplementary Figure 1C-1G). While the differences were not significant for breast, pancreatic and colorectal cancer, IDH mutant tumors still exhibit a lower LST score (Supplementary Figure 1H-1J).

The overall survival of IDH1/2 mutant tumors was also significantly higher than BRCA1/2-mutant tumors (Figure 1D). This difference may be partly explained by the more indolent nature of IDH1/2-mutated gliomas compared to BRCA1/2- mutated gliomas (Supplementary Figure 1K-1L). The lower numbers of non-glioma mutant-IDH cases precluded any detailed analysis of significant differences in survival across tumor types between IDH1/2-mutated and BRCA1/2-mutated tumors (Supplementary Figure 1M-1T).

IDH mutant tumors were not devoid of mutational signatures. In comparison to BRCA1/2-mutated and DDR-deficient tumors, IDH-mutant tumors showed increased signal for mutational signatures associated with CpG methylation (Supplementary Figure 1U). Point mutations were similarly decreased in IDH mutant tumors; APOBEC signatures and silent and non-silent mutation signatures were significantly reduced in IDH1/2-mutated compared to BRCA1/2-mutated or DDR-mutated tumors (Supplementary Figure 1V-X). This analysis of neary 50,000 cases (MSK-IMPACT database) and nearly 9,000 cases (TCGA database) does not support the hypothesis that IDH1/2-mutated tumors are tumorigenic as a result of impaired homologous recombination.

### Growth characteristics of IDH1/2-mutated cells

Previous reports have shown that cells with IDH mutations exhibit increased double-strand breaks and sensitivity to poly-ADP-ribosylation (PARP) inhibition when proliferating in culture (Sulkowski et al., 2017, 2020, 2020). To confirm and characterize this phenotype further, we generated a panel of cell lines expressing inducible IDH1 or IDH2 wild-type or mutated isoforms. Confirming functional expression of the neomorphic enzymes, inducible expression of the mutant isoforms IDH1-R132H or IDH2-R172K in the osteosarcoma cell line U2OS resulted in production of the oncometabolite 2-hydroxyglutarate (2HG) to at least 200-fold higher levels compared to wild-type (Figure 2A-B, D-E). Expression of the IDH1- or IDH2-mutant isoforms also led to significant decreases in net cell accumulation in U2OS cells (Figure 2C,F). This phenotype was also seen with the IDH1-R132H isoform in the chondrosarcoma cell line CH2879 and with the IDH2-R172K isoform in the non-transformed murine progenitor cell line C10T1/2 and human tumor cell lines HCT116, RKO and HT29) (Figure 2G).

**Figure 2.**
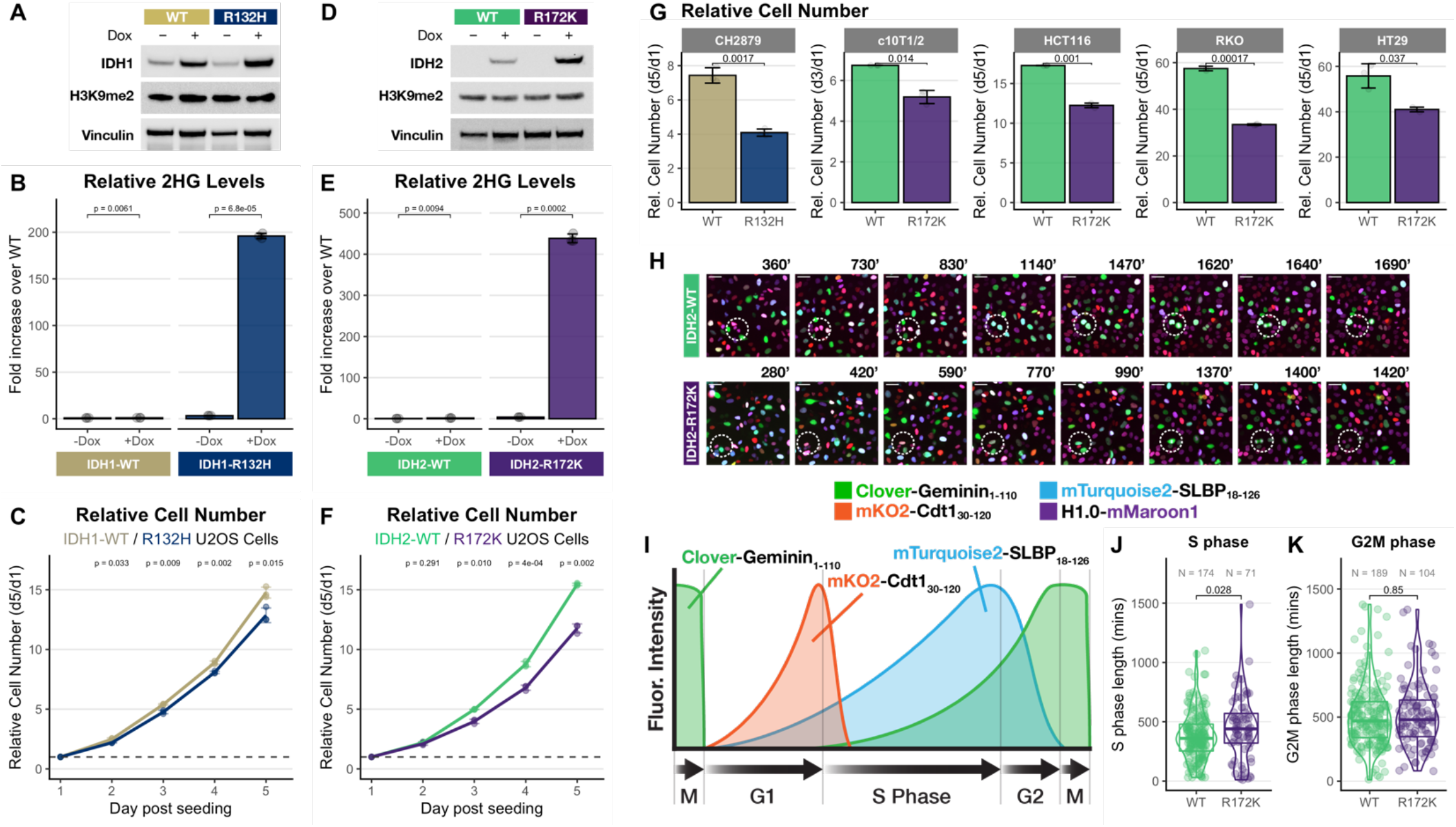
IDH1/2 mutant cells have a slower proliferation rate than wild-type cells and have a prolonged S phase. A. Western blot of lysates from IDH1-WT and IDH1-R132H transduced U2OS cells probed for indicated proteins. B. Relative 2HG levels in U2OS IDH1-WT and IDH1-R132H cells. C. Relative cell number per day after seeding in U2OS IDH1-WT and IDH1-R132H. D-F. As per A-C for U2OS IDH2-WT and IDH2-R172K cells. G. Relative cell number in indicated cell lines transduced with IDH1-WT or IDH1-R132H and IDH2-WT or IDH2-R172K. H. Example images from timelapse fluorescent microscopy in U2OS IDH2-WT and IDH2-R172K cells transduced with 4 color FUCCI system (scale bar is 50 μm). I. Cartoon illustrating cell cycle distribution of FUCCI4 system reporters. J. Quantified S phase length of U2OS IDH2-WT and IDH2-R172K cells. K. Quantified G2M phase length of U2OS IDH2-WT and IDH2-R172K cells. Error bars in B-C, E-F and G represent standard deviation from the mean of 3 replicates. P values in B-C, E-F, and G correspond to unpaired t-test. P values in J-K correspond to a Wilcoxon test.

To better understand the nature of the impaired proliferation in IDH-mutated cells, we generated IDH2-WT and IDH2-R172K U2OS cells carrying the FUCCI4 system, a set of degradable fluorescent reporters whose pattern of degradation allows estimation of the timing of each cell cycle phase (Bajar et al., 2016)(Figure 2H-I and Supplementary Figure 2A). Using this system in conjunction with time-lapse microscopy and automated cell-tracking and fluorescence quantification software, we found that IDH2-R172K mutant cells had a delay in S phase with no alteration in G2M (Figure 2J-K). This delay in IDH2-R172K transduced cells was also seen in RKO and C10T1/2 cells (Supplementary Figure 2B-E). These results show that IDH2-R172K mutations impair proliferation as a result of a prolongation of S phase.

### IDH mutations slow DNA replication in a heterochromatin-dependent manner

Since 2HG leads to enhanced methylation of marks associated with heterochromatin (Lu et al., 2012; Schvartzman et al., 2019; Sulkowski et al., 2017), we wondered whether the slower proliferation of IDH-mutant cells was a result of slower DNA replication through areas of heterochromatin. To assess the rate of DNA replication in a per-cell manner we quantified the incorporation of the thymidine analogue EdU after a 20 minute pulse of EdU was added to asynchronously growing cells. In this assay, cells are harvested and fixed after this limited EdU pulse, and click chemistry is used to fluorescently label EdU. Using this assay, we found that the labeling of EdU positive cells in IDH-mutated cells was significantly reduced compared to IDH-WT cells (Figure 3A and Supplementary Figure 3A-C) with only minor changes in the proportion of cells in each cell-cycle phase (Figure 3B). This effect was not due to differing amounts of DNA content per cell (e.g. as a result of genomic instability). A 2 hour pulse of cell-permeable 2HG also showed a reduction in the EdU-positive peak (Figure 3C). These effects were also seen for the colorectal cancer cell line RKO and for the non-transformed murine mesenchymal progenitor cell line C10T1/2 (Supplementary Figures 3D-L and 3M-N). Addition of bleomycin at doses known to induce double-strand-breaks led to no changes in the intensity of EdU signal (Supplementary Figure 3O-Q).

**Figure 3.**
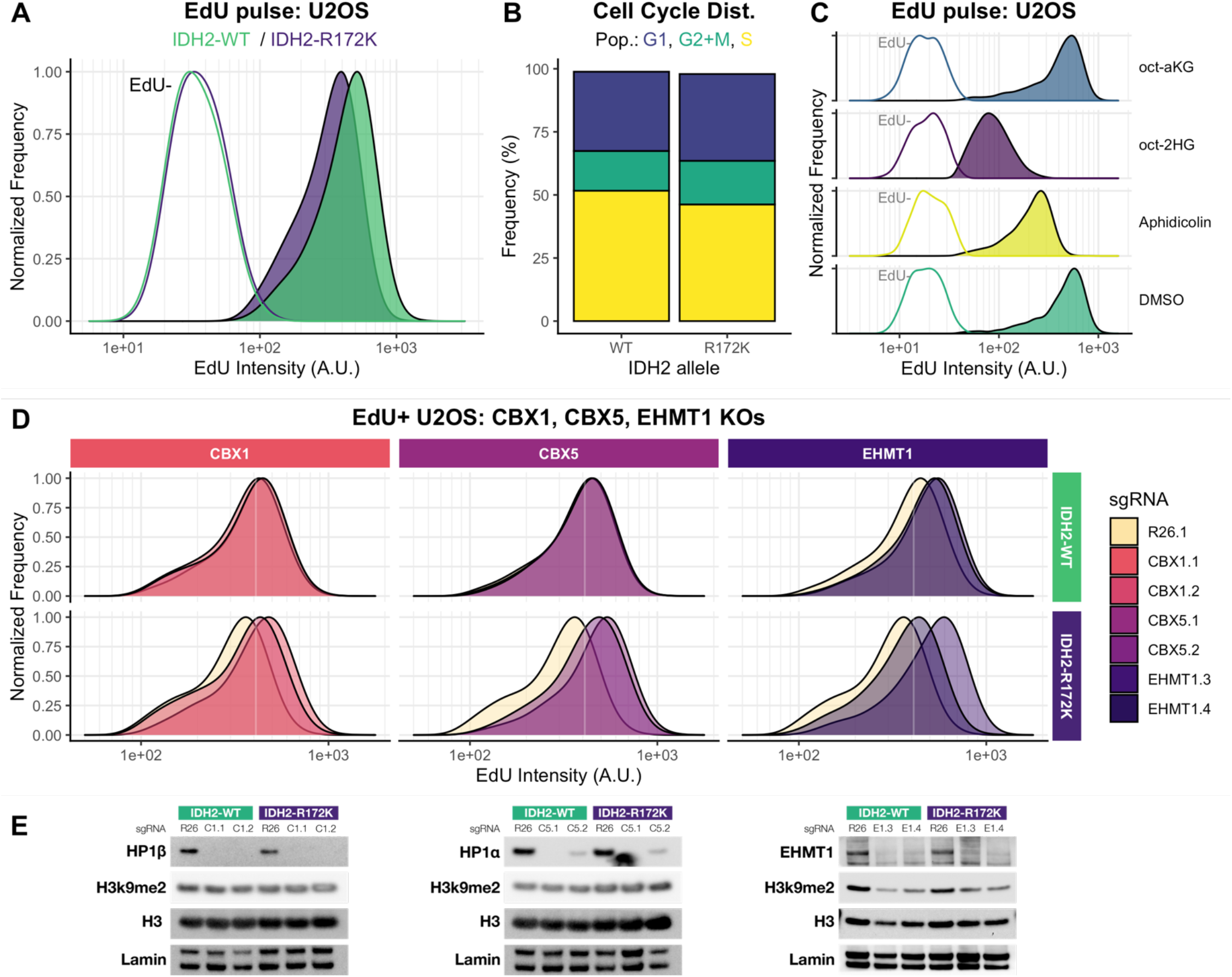
IDH2 mutations and 2HG slow DNA replication in a heterochromatin-dependent manner. A. Distribution of EdU signal intensity in EdU+ (solid) and EdU- (hollow) IDH2-WT and IDH2-R172K U2OS cells. B. Cell cycle distribution frequencies in U2OS IDH2-WT and IDH2-R172K cells. C. EdU signal intensity distribution in U2OS parental cells treated with indicated drugs for 2 hours. D. EdU+ signal intensity distribution in IDH2-WT and IDH2-R172K cells co-transduced with CRISPR-Cas9 vectors targeting Rosa26 (yellow fill) and CBX1 (HP1β), CBX5 (HP1α) and EHMT1 (GLP). E. Western blot of lysates from cell lines used in D probed for indicated proteins.

To determine whether the observed slowing of DNA replication was a result of increased heterochromatin in IDH2-mutant cells, we generated CRISPR-Cas9 knock-outs of three genes encoding proteins required for heterochromatin function (CBX1 and CBX5 genes coding for the heterochromatin proteins HP1β and HP1 a, respectively; and EHMT1 coding for the H3K9 methyl transferase GLP (Ayyanathan et al., 2003; Chen et al., 2012; Eissenberg et al., 1990). Loss of HP1β, HP1α or EHMT1 in IDH2-R172K cells led to a level of EdU incorporation similar to that seen in IDH2-WT cells (Figure 3D-E). Knock-out of CBX1 or CBX5 in IDH2-WT cells did not result in further increases in EdU incorporation while knock-out of EHMT1 showed a modest increase in EdU incorporation. These results demonstrate that IDH2-mutation results in slowing of DNA replication and that this phenotype correlates with increased heterochromatin.

### Slower global DNA replication in IDH-mutant cells is a result of individual replication fork slowing

The finding of impaired EdU incorporation after a 20 minute pulse of EdU in IDH-mutant cells is consistent with reduced DNA replication. This could be either a result of an altered pattern of active replication origins (e.g. fewer active origins in IDH-mutant cells) or slower individual replication fork progression of some or all replication forks in IDH-mutant cells. To determine whether the replication origin landscape was altered as a result of oncogenic IDH-mutations, we performed Repli-SEQ in IDH2-WT and IDH2-R172K U2OS cells. In Repli-SEQ, asynchronously growing cells are treated with the thymidine analogue BrdU for 2 hours and subsequently harvested and sorted into early replicating or late replicating according to DNA content. Early and Late replicating pools are then used to generate DNA libraries for next-generation sequencing (Marchal et al., 2018, 2018; Pope et al., 2014). An analysis of the ratio of early to late replicating regions in 50kb windows showed nearly identical signals between IDH2-WT and IDH2-R172K U2OS cells (Figure 4A-B) and RKO cells (Supplementary Figure 4).

**Figure 4.**
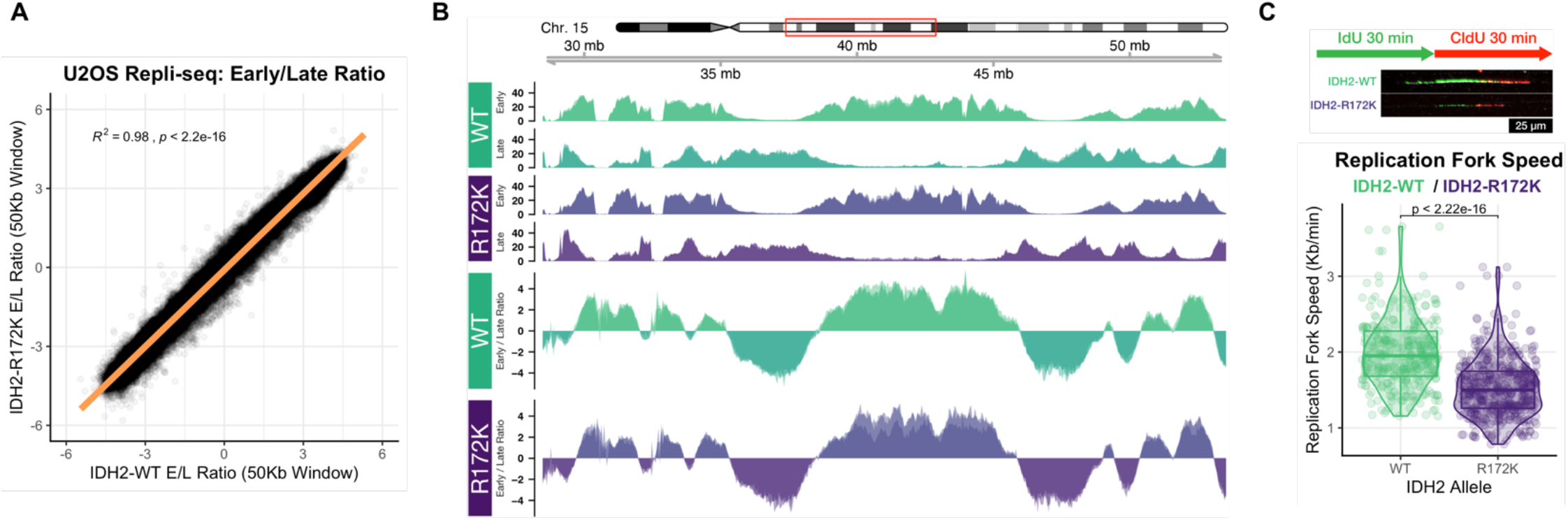
IDH-mutations lead to slower replication fork progression with minor changes in the replication landscape. A. U2OS IDH2-WT and IDH2-R172K cells were incubated asynchronously in BrdU for 2 hours and sorted by DNA content into early (E) and late (L) S phase fractions. Following fragmentation, BrdU immunoprecipitation and repli-seq library generation were carried out. Shown is the distribution of E/L ratios of reads per 50 Kb windows by IDH status (IDH2-WT, horizontal axis; IDH2-R172K, vertical axis). A downwards shift indicates areas that replicated earlier in IDH2-WT cells while a shift upwards indicates areas that replicated earlier in IDH2-R172K cells. B. Early and Late repli-seq tracks from IDH2-WT and IDH2-R172K U2OS cells. Ratio of early/late tracks plotted in lower two sections. C. Top: cartoon and example immunofluorescence for DNA combing assay. Bottom: quantification of replication fork speed derived from combined IdU and CldU tracks from IDH2-WT/R172K U2OS cells. Scale bar is 25 μm. In A, R^2^ and p value correspond to the Pearson correlation coefficient. P value In C corresponds to a Wilcoxon test.

To test whether IDH-mutation resulted in slower individual DNA replication forks, we performed DNA combing and immunofluorescence for thymidine analogues after serial pulses of IdU and CldU in asynchronously growing IDH2-WT and IDH2-R172K cells (Michalet et al., 1997; Técher et al., 2013). Analysis of individual forks showed a significant decrease in the median replication fork speed of forks obtained from IDH2-R172K cells compared to IDH2-WT cells (Figure 4C).

### S-phase specific double strand breaks in IDH mutant cells result from heterochromatin dependent replication stress and require poly-ADP ribosylation for correct processing

Prior work had shown that IDH-mutant cells incur a higher number of doublestrand breaks as seen by comet assays and immunofluorescence with pH2AX (Sulkowski et al., 2017, 2020, 2020). Since we had observed a prolongation of S phase and slower replication fork progression in IDH2-R172K-expressing cells, we wondered whether this DNA replication defect corresponded with a higher number of replication-related DNA breaks. To map whether pH2AX foci had any relation to DNA replication, we performed immunofluorescence for pH2AX in conjunction with a 30 minute pulse label of EdU, thus distinguishing actively proliferating cells from cells in either G1 (2n DNA content) or G2/M (4n DNA content).

Untransformed C10T1/2 mesenchymal progenitors expressing IDH2-R172K had a significantly higher amount of pH2AX foci that was enriched in EdU-positive cells (Figure 5A-C and Supplementary Figure 5A-B). This correlation was also observed in the tumor cell line U2OS transfected with IDH2-R172K-expressing vectors, where very little signal was observed in EdU-negative cells (Figure 5D-F) and in IDH2-R172K-expressing RKO cells (Supplementary Figure C-D). The increase in pH2AX signal in EdU-positive cells was dependent on heterochromatin, as loss of CBX1 (HP1β), CBX5 (HP1α) and EHMT1 (GLP) resulted in a reduction of the pH2AX signal to wild-type levels (Figure 5G and Supplementary Figure 5E).

**Figure 5.**
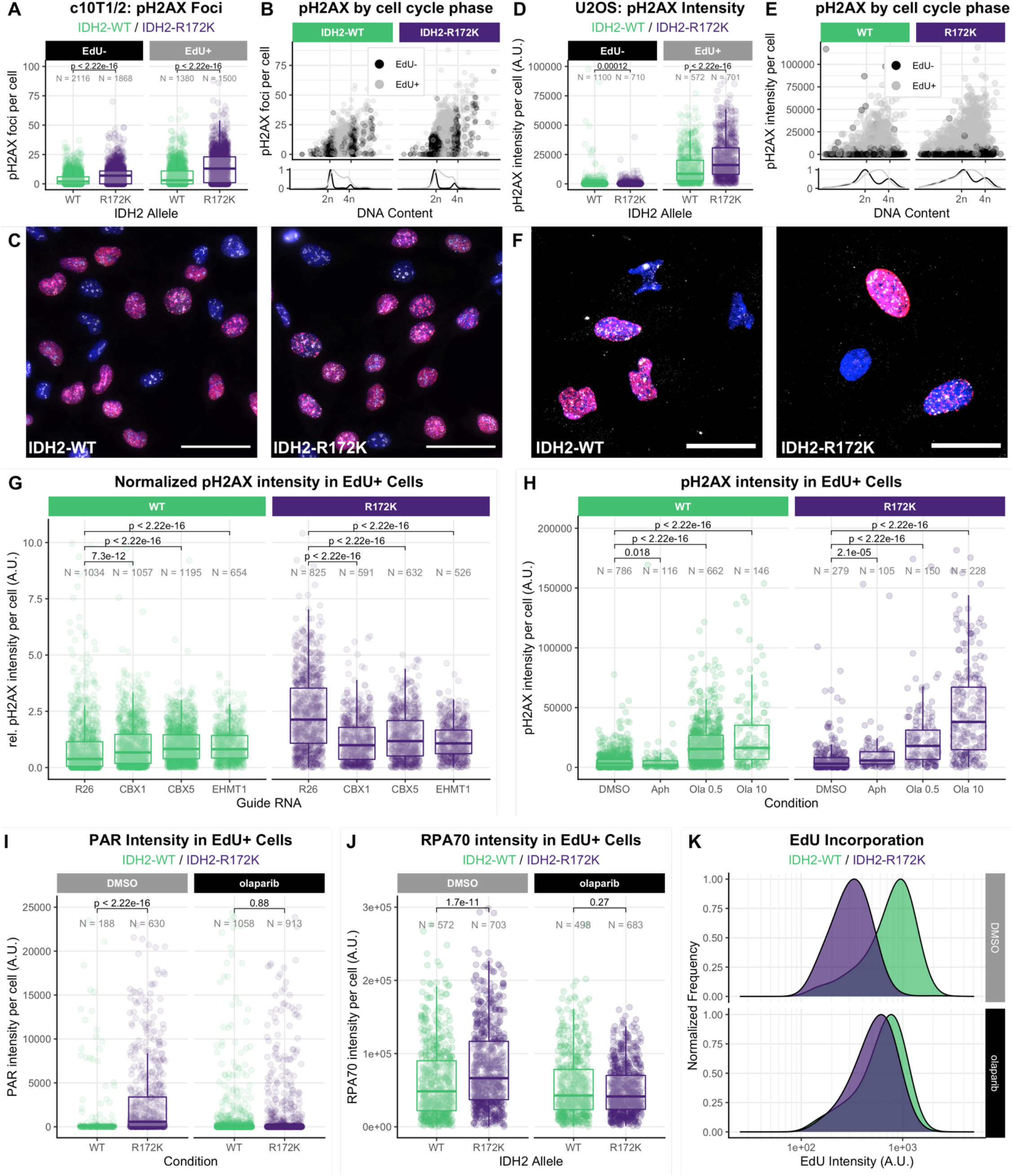
The increase in DNA double-strand breaks in IDH mutant cells is associated with DNA replication stress and chromatin PAR-ylation. A. Distribution of pH2AX foci in C10T1/2 IDH2-WT and IDH2-R172K cells by EdU signal intensity. B. pH2AX foci quantification from A plotted by EdU signal intensity (EdU-in black, EdU+ in grey) and DNA content determined by quantification of DAPI signal on horizontal axis. Density distribution of DNA content shown in lower panel. C. Example immunofluorescence images of IDH2-WT and IDH2-R172K C10T1/2 cells; DAPI in blue, EdU in magenta, pH2AX in white. Scale bar is 50 μm. D. Distribution of pH2AX signal in U2OS IDH2-WT and IDH2-R172K cells by EdU signal intensity. E. pH2AX signal from D by EdU signal intensity (EdU-in black, EdU+ in grey) by DNA content determined by quantification of DAPI signal on horizontal axis. Density distribution of DNA content shown in lower panel. F. Example immunofluorescence images of IDH2-WT and IDH2-R172K U2OS cells; DAPI in blue, EdU in magenta, pH2AX in white. Scale bar is 25 μm. G. Relative pH2AX signal intensity in U2OS IDH2-WT/R172K cells transduced with indicated CRISPR/Cas9 vectors for the indicated genes. H. Relative pH2AX signal intensity in U2OS IDH2-WT and IDH2-R172K cells treated with indicated drugs. I. PAR fluorescence intensity in U2OS IDH2-WT and IDH2-R172K cells treated with vehicle (DMSO) or olaparib. J. RPA70 fluorescence intensity in U2OS IDH2-WT and IDH2-R172K cells treated with vehicle (DMSO) or olaparib. K. EdU+ signal intensity distribution for U2OS IDH2-WT and IDH2-R172K cells treated with vehicle (DMSO) or olaparib. P values in A-B, D-E and G-J correspond to pair-wise Wilcoxon tests. In G, Kruskal-Wallis P values are 7e-32 (IDH2-WT) and 3.6e-66 (IDH2-R172K). In H, Kruskal-Wallis P values are 1.2e-110 (IDH2-WT) and 2.1e-54 (IDH2-R172K).

IDH-mutant cells have been shown to be more sensitive to inhibition of poly-ADP ribose polymerase. We wondered whether PARP-inhibition had an effect at the level of DNA replication stress observed in IDH mutant cells. We found that pH2AX intensity was significantly higher in EdU+ U2OS cells treated with the PARP-inhibitor olaparib (Figure 5H), and this effect was greater in IDH2-R172K cells compared to WT. The poly-ADP-ribosylation (PAR) signal was significantly higher in EdU+ IDH2-R172K cells, and consistently decreased upon treatment with olaparib (Figure 5I). Olaparib also resulted in a decrease in the overall intensity of the single-stranded DNA (ssDNA) binding protein RPA70, suggesting that IDH2-R172K cells incurred a higher amount of replication stress in S phase and that PAR-ylation was required for recruitment of RPA70 to ssDNA (Figure 5J). Consistent with this possibility, we found that olaparib treatment rescued the decrease in EdU incorporation rate observed in IDH2-R172K cells (Figure 5K).

## Discussion

A growing number of intracellular metabolites has been linked to the enzymatic modification of DNA and chromatin-associated proteins. Whether variations in these intracellular metabolites impact chromatin in a manner that affects the integrity of DNA replication remains unclear. The above results suggest that IDH-mutant cells have increased heterochromatin resulting in slower DNA replication and enhanced levels of heterochromatin-associated replicative stress during cell division. Poly-ADP-ribosylation appears to be required to process and repair these lesions, as addition of the PARP inhibitor olaparib results in further increases in pH2AX signal and double strand breaks.

Our work suggests that alpha-ketoglutarate-dependent dioxygenases play a role in permitting faithful and processive replication of heterochromatin. This is highlighted by the fact that 2HG-dependent DNA-breaks and replication slowing are rescued by mutations in essential components of heterochromatin (HP1β, HP1α and EHMT1).

The analysis of over 50,000 patient samples shows that IDH mutations do not confer a phenotype of genomic instability, as seen in tumors harboring mutations in the BRCA1/2 tumor suppressors or with loss-of-function mutations in other DNA-Damage-Response genes. This argues that IDH mutations do not result in functional impairment of homologous recombination-mediated repair. However, prior studies have shown a synthetic lethality of IDH-mutations with inhibition of poly-ADP-ribose polymerase activity (Gbyli et al., 2022; Lu et al., 2017; Sulkowski et al., 2017, 2020, 2020). Recent work has shown that the mechanistic link between loss of BRCA1/2 protein function and sensitivity to PARP inhibitors is a result of DNA replication gaps (Cantor, 2021; Cong et al., 2021; Genois et al., 2021). In contrast, IDH-mutant cells display increased amounts of heterochromatin and S-phase-specific replication gaps that can sensitize cells to PARP inhibition in the presence of normal homologous recombination. Clearer identification of where the sites of replicative stress are located relative to the heterochromatin regions will require analysis of single replication fork dynamics with sequence mapping resolution, a technique that has not yet been feasible in mammalian cells (Claussin et al., 2022).

A further question that arises from our work is whether a 2HG-dependent-like process occurs in physiological settings outside that of IDH-mutant tumors. Sulkowski et al (Sulkowski et al., 2017, 2020, 2020) identified the alphaketoglutarate-dependent dioxygenases responsible for the 2HG-dependent double strand breaks (Kdm4a and Kdm4b). These KDMs hydroxylate H3K9 di- and tri-methyl marks, ultimately leading to their removal. Of the multiple H3K9 demethylases, Kdm4b has one of the highest Michaelis-Menten constants (Km) for oxygen (Chakraborty et al., 2019), marking it as a potential oxygen sensor for chromatin. These findings suggest that tissues and tumors with lower oxygen concentrations (~ 5-20 μM Oxygen) may be susceptible to increased amounts of replicative stress (Ast and Mootha, 2019) and thus sensitive to PARP inhibition.

## Author Contributions

Conceptualization, J.M.S., and C.B.T.; investigation and methodology, J.M.S., G.F.; formal analysis, validation and curation, J.M.S., G.F., H.W., W.C., A.S., K.M., F.S.V., R.K.; writing – original draft, J.M.S.; writing – review & editing, J.M.S. and C.B.T.; visualization, J.M.S.; funding acquisition, J.M.S. and C.B.T.

## Acknowledgements

We thank members of the Thompson lab for insightful discussions during manuscript preparation. We thank Iestyn Whitehouse and the Integrative Genomics Operation at MSKCC for experimental advice. We thank Eric Chan from the MSKCC Molecular Cytology Core for assistance with microscopy analysis. J.M.S. was a Hope Funds for Cancer Research Fellow, a recipient of a Pittsburgh Cure Sarcoma research award and received funding from a Department of Defense, Congressionally Directed Medical Research Program Career Development Award (W81XWH-20-PRCRP-CDA). This work is supported by grants from the NCI (to C.B.T.), by the cancer center support grant (P30 CA008748) to Memorial Sloan Kettering Cancer Center and by the Marie-Josée and Henry R. Kravis Center for Molecular Oncology and the National Cancer Institute Cancer Center Core Grant No. P30-CA008748. We gratefully acknowledge the members of the Molecular Diagnostics Service in the Department of Pathology at Memorial Sloan Kettering Cancer Center.

## Methods

### Tumor mutational profiling and LST scores

Solid tumors from ~50,000 patients treated at MSKCC were sequenced using MSK-IMPACT, a capture-based next-generation sequencing platform that can detect mutations, copy number alterations, and select rearrangements in 341-505 cancer-associated genes, depending on the version of the panel. The MSK-IMPACT assay achieves high depth of sequencing (800x) and is performed in a Clinical Laboratory Improvement Amendments (CLIA)–certified molecular laboratory, as previously described (Cheng et al., 2014). Matched blood specimens were used to filter out germline alterations. Somatic mutations, copy number alterations, and gene fusions were called for each sample using the MSK-IMPACT computational pipeline, as previously published (Zehir et al., 2017).

The FACETS algorithm was used to infer allele-specific copy number and estimate tumor purity for MSK-IMPACT samples (Shen and Seshan, 2016). The FACETS Suite package was used to generate large-scale state transition (LST) scores, which are the number of chromosomal breaks between regions that are adjacent and are at least 10 Mb in size, from the copy-number data provided by FACETS (https://github.com/mskcc/facets-suite). LST scores for TCGA samples were calculated using methods by Popova et al. (Popova et al., 2012) and were obtained from the work published by the TCGA Pan-Cancer analysis of DNA Damage Repair Deficiency working group (Knijnenburg et al., 2018).

### Survival Plots

Survival plots were generated from MSK-IMPACT and TCGA data using R language with the survminer (https://github.com/kassambara/survminer) and survival (https://github.com/cran/survival) packages.

### Cell culture

U2OS, RKO, C10T1/2 (C3H/10T1/2, Clone 8), HT29, HCT116 cells were obtained from ATCC. CH2879 cells were obtained from SKI. All cells were cultured at 37°C in 5% CO2 in high glucose DMEM supplemented with 10% FBS. All lines were determined to be mycoplasma free using MycoAlert Mycoplasma Detection Kit (Lonza). For growth assays, 25-50K cells (depending on cell line) were seeded in 12-well tissue culture plates on day 0 in triplicate. Cells were counted using a Beckman-Coulter Multisizer 3 on days 1 and subsequent time-points as indicated. IDH2-WT and IDH2-R172K inducible vectors were characterized previously (Schvartzman et al., 2019). Retroviral particles were produced in 293T cells by transfecting pCG-gag-pol and pCMV-VSV-G packaging plasmids (Addgene) together with the corresponding retroviral plasmid. For CRISPR knock-out lines, pLentiCRISPRv2 plasmids were cloned with indicated sgRNA sequences and used to transfect 293T cells together with pMD2 and psPAX2 vectors for lentiviral generation. Viral supernatants (retroviral or lentiviral) were harvested 48 hours after transfection, filtered through 0.45 μm nylon filters and transduced into target cells in the presence of 8 μg/ml polybrene.

### GC-MS analysis of 2-hydroxyglutarate

Cells were plated on 6 well plates in triplicate at a density of 400e3 cells/well the day before harvesting. Media was aspirated and cells were fixed with 80% methanol containing 20 μM deuterated 2HG as an internal standard (D-hydroxyglutaric- 2,3,3,4,4-d5) pre-chilled to −80°C for 30 minutes at −80°C). Lysates were then incubated for 4 hours at −80°C, scraped, transferred to 1.5 mL tubes and centrifuged at 20,000 g for 20 minutes at 4°C. The supernatant was dried by spin vacuum and then resuspended in 40 mg/mL methoxyamine in pyridine. Each sample was then derivatized with MSTFA with 1% TMCS for 30 minutes at 30°C. One microliter of trimethylsilyl-derivatized organic acids was analyzed using an Agilent 7890A GC equipped with an HP-5MS capillary column and connected to an Agilent 5975 C mass selective detector operating in splitless mode with electron impact ionization. Relative quantification of 2HG was performed from extracted ion chromatograms for 2HG (m/z: 349) normalized to the D5-2HG internal standard (m/z: 354) and corrected by protein concentration per sample.

### Western blotting

Protein extracts were prepared using 1X RIPA buffer with protease and phosphatase inhibitors (Thermo # 1860932 and Thermo # 78428, respectively). Equal amount of protein was separated on NuPAGE Bis-Tris gels (Life Technologies) and transferred to nitrocellulose membranes. Membranes were blocked in 5% milk in TBST and incubated with indicated primary antibodies. After secondary antibody incubation and 1-shot ECL solution (Kindle Biosciences LLC) incubation, blots were scanned using a Biorad ChemiDoc Touch Imaging System.

### Time-lapse microscopy and analysis

Cells were transduced with FUCCI4 viral vectors (Bajar et al., 2016) coding for Clover– Geminin_1–110_, mKO2–Cdt1_30–120_ (plasmid/virus 1) and mTurquoise2–SLBP_18–126_ and H1.0–mMaroon1 (plasmid/plasmid 2). Cells positive for markers from both vectors were sorted using a Sony SH800S cell sorter and seeded on μ-Slide 8 Well high Glass Bottom chambered slides (Ibidi). Time-lapse images were acquired on a Zeiss ZEN imaging system.

Nuclei in time lapse images were tracked and all channel expression levels quantitated with the Fiji plugin Trackmate (sigma=12.5px, threshold=30, max motion= 30px, max motion gap=15px). Tracks were imported into MATLAB for analysis. Custom scripts filtered cells automatedly for appropriate cells in which to quantify cell cycle phase lengths. Expression over time in each channel was smoothed with a moving median filter of width 15 frames. Maxima and minima were found in mKO2-Cdt1 and mTurquoise2-SLBP channels. The location of maximal drop in signal was found in the Clover-Geminin channel. This drop in Clover-Geminin was considered significant (indicating it captured a genuine division) if it reached a threshold of 900 arbitrary units. Cells without significant expression variation in mKO2-Cdt1 and mTurquoise2-SLBP (between the global maxima and minima found) over time were discarded based on a threshold of 300 arbitrary units. Also discarded were cells with mKO2-Cdt1 or mTurquoise2-SLBP instantaneous changes over 300 arbitrary units (as sudden changes this large are not expected, even at division, and so change indicates a likely tracking error).

G1 was measured as the time period between a Clover-Geminin drop and mKO2-Cdt1 max. S was measured as the period between mKO2-Cdt1 max and mTurquoise2-SLBP max. G2+M were measured as the difference between mTurquose2-SLBP max and Clover-Geminin drop.

Each cell cycle phase was measured in the set of tracks which passed checks for significant change in the relevant channels, length >160 frames and in which the two features (maxima or maximal instantaneous change) were found in the correct order indicating for that specific cell the movie captured the entirety of the desired cell cycle.

Slope of Clover-Geminin during S phase in arbitrary units per minute was estimated for each valid S phase cell via linear fit to the Clover-Geminin signal between mKO2-Cdt1 and mTurquoise2-SLBP peaks.

### EdU incorporation and quantification by flow cytometry

Asynchronously growing cells were pulsed with 20 μM EdU for 20 minutes and harvested by trypsinization into cold PBS. Cells were fixed in 4% formaldehyde in PBS for 15 minutes and permeabilized with 0.05% Triton-X in PBS for 15 minutes. EdU was labelled using click chemistry using sulfo-Cyanine 3 Azide (Lumiprobe), CuSO_4_ and Sodium Ascorbate. Cells were then incubated in Hoechst 33342 with RNAse A in PBS with 1% FBS and EdU and DNA content (Hoechst 33342) was detected using a BD LSRFortessa Cell Analyzer.

### Repli-seq

Repli-seq was carried out as previously described (Marchal et al., 2018, 2018). Briefly, asynchronously growing cells were pulsed with 100 μM BrdU for 2 hours and harvested by trypsinization. Cells were then resuspended in ice-cold PBS and fixed with ice-cold EtOH to a concentration of 75%. Fixed cells were labelled for DNA content with PBS/FBS/Propidium Iodide/RNAse A solution and sorted for early and late replicated DNA by PI-content. Cells were then lysed and genomic DNA extracted using Zymo Quick-DNA Microprep Kit. DNA was fragmented using a Covaris E220 ultrasonicator and genomic libraries generated using the NEBNext Ultra DNA Library Prep Kit. Fragments were then immunoprecipitated with primary mouse anti-BrdU and secondary rabbit anti-mouse antibodies. After a cleaning step, libraries were amplified with NEBNext Multiplex Oligos for Illumina. Libraries were size-selected with AmpureXP beads and quality control performed with Tapestation (Agilent), Q-bit (ThermoFisher) and KAPA quantification (Roche). Sequencing was carried out in an Illumina NextSeq 500 platform using paired-end 100 bp sequencing.

For analysis, sequencing reads were trimmed and filtered for quality (Q >= 15) and adapter content using version 0.4.5 of TrimGalore (https://www.bioinformatics.babraham.ac.uk/projects/trim_galore) with version 1.15 of cutadapt and version 0.11.5 of FastQC. Reads were aligned to human assembly hg19 with version 2.3.4.1 of bowtie2 (http://bowtie-bio.sourceforge.net/bowtie2/index.shtml) and were deduplicated using MarkDuplicates in version 2.16.0 of Picard Tools. The BEDTools suite (http://bedtools.readthedocs.io) and R was used to process the bam files as described in steps 80-92 for paired-end data in (Marchal et al., 2018, 2018). Briefly, read density was compiled over 50 kb sliding windows for both Early and Late and the final replication timing value was calculated as log2(Early:Late ratio) and adjusted via quantile normalization in R.

### DNA combing and fiber analysis

Cells were grown asynchronously and pulsed with 100 μM IdU for 30 minutes followed by 2 washes in pre-warmed media. Cells were then pulsed with 100 μM CldU for 30 minutes and washed in ice-cold PBS followed by trypsinization and resuspension in ice-cold media. Cells were then counted and 125,000 cells were used for DNA plugs by mixing cells in 45 μL of resuspension buffer at 50°C and 2% low-melting agarose at 50°C. Plugs were formed in dedicated wells (Genomic Vision) and incubated at 4°C for 30 minutes. Plugs were then transferred into proteinase K buffer (10 mM Tris pH 7.5, 100 mM EDTA pH 8, 0.1% N-lauroylsarcosine, 0.2% Na-deoxycholate (Genomic Vision) with 1 mg/ml proteinase K and incubated at 50°C overnight. The following day, plugs were washed 3 times in wash buffer (100 mM Tris pH 7.5, 10 mM EDTA pH 8, 1M NaCl, Genomic Vision) for 1 hour each and then transferred to round bottom micro-centrifuge tubes containing 50 mM MES monohydrate. Plugs were incubated at 68°C for 20 minutes and then at 42°C for 10 minutes and agarase was added to dissolve the agarose plug. DNA plugs were incubated at 42°C overnight and poured into dedicated reservoirs for DNA combing (Genomic Vision). Combing was performed using a Genomic Vision Combing Instrument on dedicated coverslips. Coverslips were then baked at 65°C for 2 hours and denatured in 0.5M NaOH, 1M NaCl for 8 minutes at room temperature followed by dehydration in 70, 90 and 100% ethanol series. Coverslips were then blocked in 5% FBS in PBS and incubated in mouse anti-BrdU and rat anti-BrdU primary antibodies in 3% FBS in PBS overnight at 4°C. Following 3 washes in 5% FBS in PBS, coverslips were incubated with anti-mouse Alexa488 and anti-rat Alexa594 secondary antibodies for 30 minutes at 37°C. For ssDNA background detection, coverslips were incubated in mouse anti-ssDNA primary antibody for 2 hours at room temperature, washed, and incubated in antimouse Alexa647 secondary antibody for 30 minutes at 37°C. Coverslips were then washed in 5% FBS in PBS and twice in PBS and then mounted with Southern Biotech Fluoromount-G without DAPI onto standard microscopy slides. Images were acquired on a Delta-Vision microscope with 10% stitching and analyzed using FiJi software to manually measure track length in blinded fashion.

### Immunofluorescence

Cells were grown on Millicell EZ 8-well glass slides (Millipore) and treated with indicated conditions. 30 minutes prior to fixation, EdU was added to a final concentration of 20 μM. After 30 minutes, cells were washed with ice cold PBS twice and fixed with 4% formaldehyde in PBS for 15 minutes at room temperature. For anti-PAR staining, cells were fixed in ice-cold Methanol for 15 minutes. Cells were then washed twice in PBST (0.01% Tween-20 in PBS) and permeabilized with 0.5% Triton-X-100 in PBS for 15 minutes at room temperature. EdU was labelled using click chemistry using sulfo-Cyanine 3 Azide (Lumiprobe), CuSO_4_ and Sodium Ascorbate for 30 minutes at room temperature in a dark environment. Cells were then washed twice with PBS and blocked in PBG solution (0.2% w/v cold water fish gelatin and 0.5% w/v BSA in PBS) overnight at 4°C or 2 hours at room temperature. Cells were then incubated in indicated primary in PBG with 0.1% Tween-20 overnight at 4°C. Secondary incubation was carried out in PBS with 4% normal goat serum for 1 hour at room temperature. Cells were then washed and nuclei counterstained with DAPI prior to mounting. Slides were imaged using a Pannoramic Confocal (3DHISTECH) and z-stack maximal projections analyzed using Fiji/ImageJ software for quantification of fluorescent intensity and area.

---

**Supplementary Figure 1.**
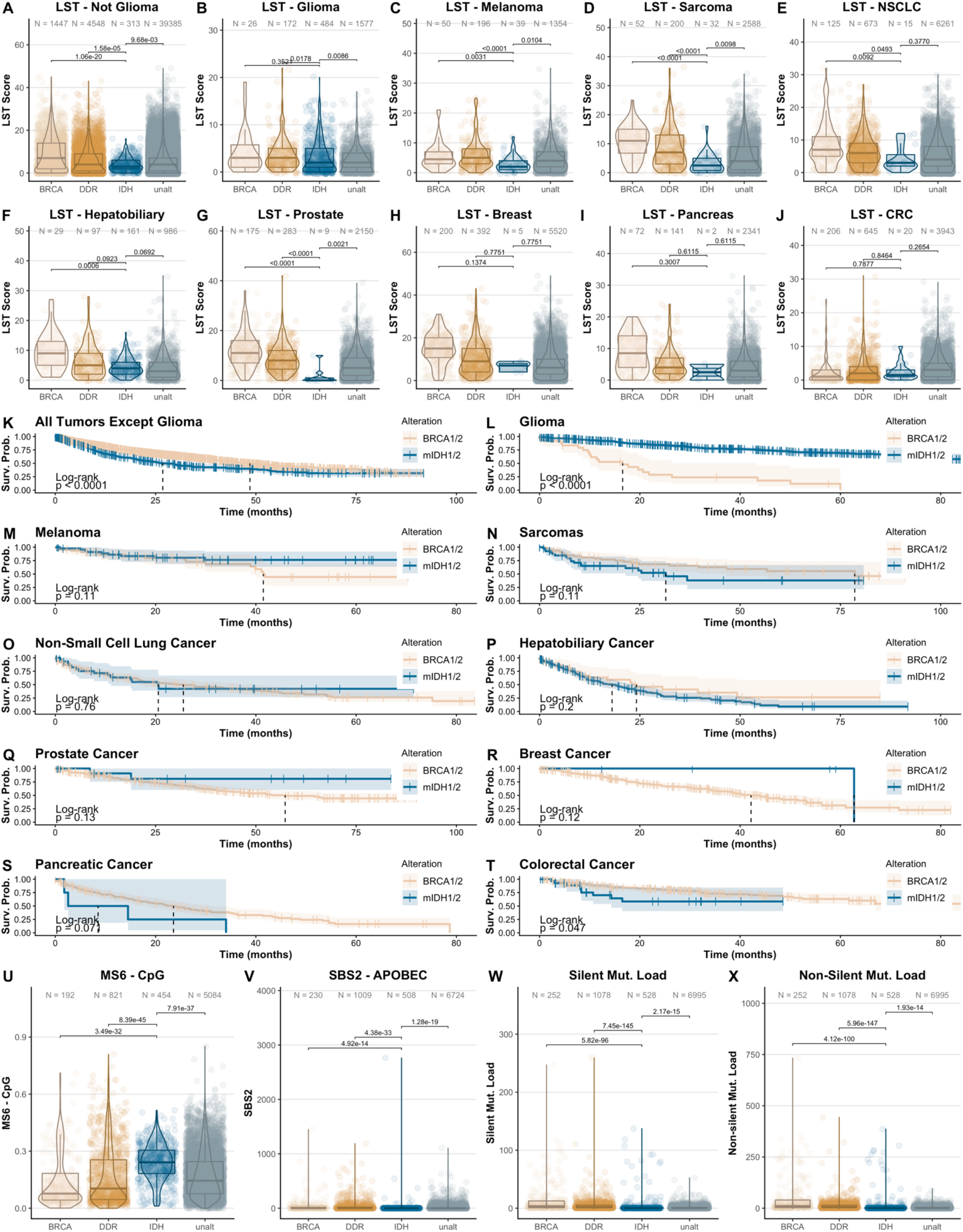
A-J. LST Scores for indicated tumor types in MSK-IMPACT data set classified by mutation status. K-T. Survival probability (Surv.Prob.) plots for indicated tumor types in MSK-IMPACT data set classified by mutation status. U-X. Indicated mutation signatures for all tumor types in TCGA data set. U: MS6 CpG mutation signature. V: SBS2 APOBEC mutation signature. W: Silent mutation load signature. X: Non-silent mutation load signature. P values for A-J and U-X correspond to a pair-wise Wilcoxon test. P values for A-J and U-X correspond to Dunn’s test after a Kruskal-Wallis. P values for Kruskal-Wallis: A) 6.2e-70, B) 0.000000079, C) 0.0000086, D) 4.3e-16, E) 1.3e-25, F) 0.000000011, G) 2.4e-39, H) 4e-58, I) 2.4e-12, J) 3.7e-26, U) 3e-52, V) 2.1e-33, W) 1.00e-297, X) 3.40e-311. P values for K-T correspond to a log-rank test.

**Supplementary Figure 2.**
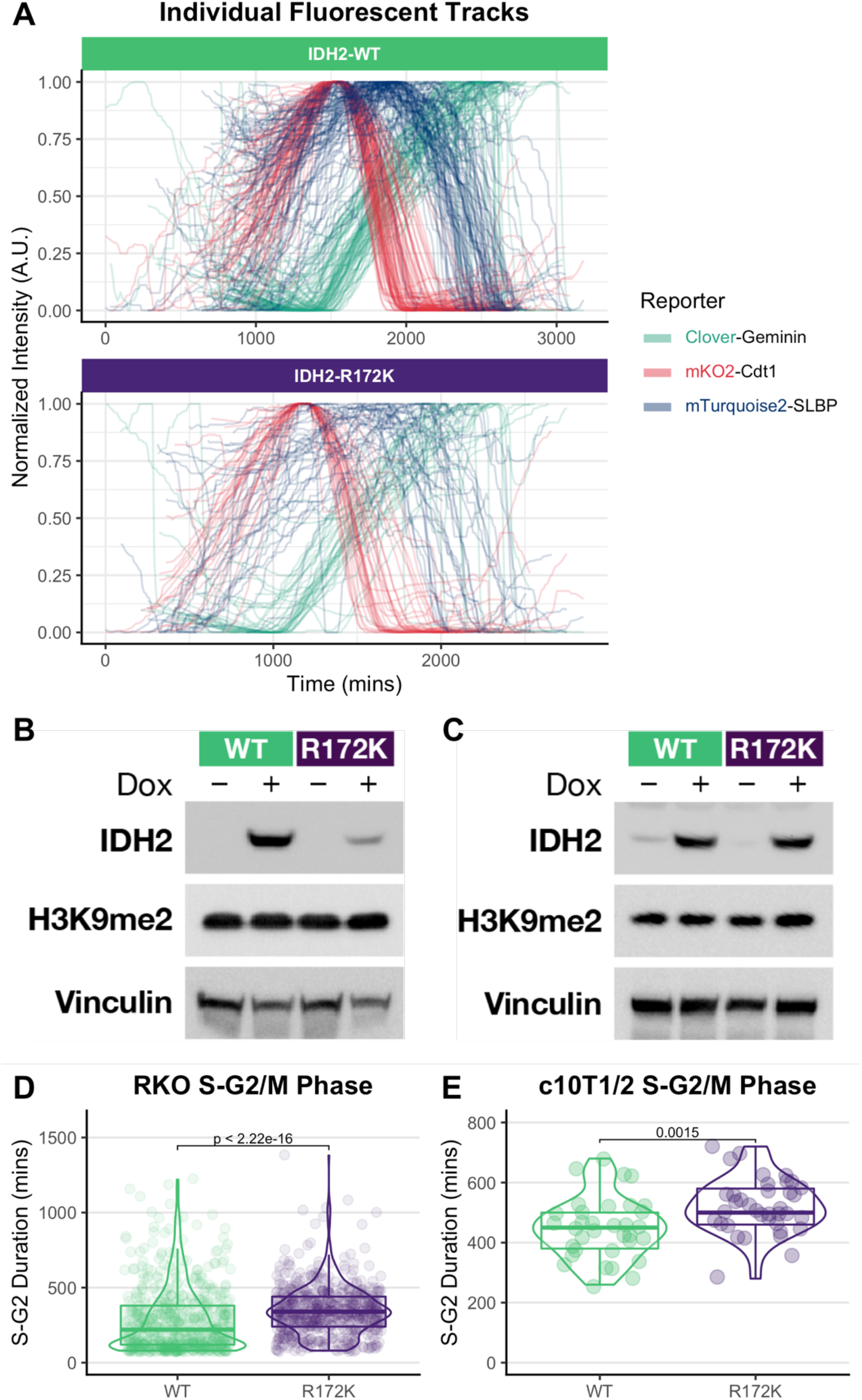
A. Overlay of normalized quantification of reporter intensity for indicated reporters in U2OS IDH2-WT and IDH2-R172 cells. B. Western blot of lysates from IDH2-WT and IDH2-R172K RKO cells probed for indicated proteins. C. Western blot of lysates from IDH2-WT and IDH2-R172K C10T1/2 cells probed for indicated proteins. D. S-G2/M length in IDH2-WT and IDH2-R172K RKO cells. E. S-G2/M length in IDH2-WT and IDH2-R172K C10T1/2 mesenchymal progenitor. P values in D-E correspond to a Wilcoxon test.

**Supplementary Figure 3.**
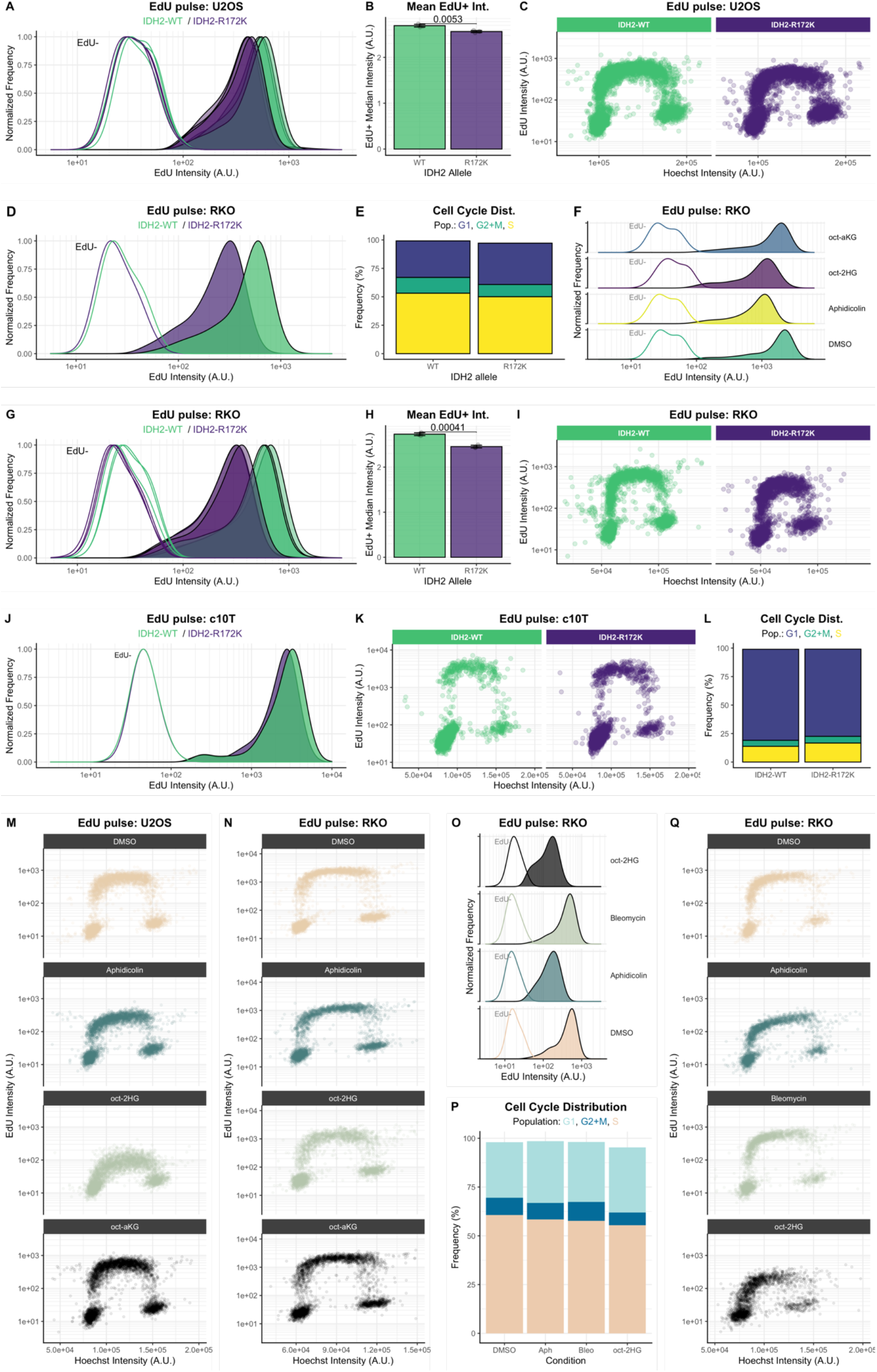
A. EdU signal intensity distribution of individual replicates for U2OS IDH2-WT and IDH2-R172K cells. B. Quantification of Mean EdU+ signal intensity from A (3 replicates per condition). C. 2-dimensional plot of EdU signal intensity for IDH2-WT and IDH2-R172K U2OS cells with DNA content (Hoechst) on horizontal axis. D. Distribution of EdU signal intensity in EdU+ (solid) and EdU- (hollow) RKO cells transduced with IDH2-WT or IDH2-R172K expressing vectors. E. Cell cycle distribution frequencies in RKO IDH2-WT and IDH2-R172K cells. F. EdU signal intensity distribution in RKO parental cells treated with indicated drugs for 2 hours. G. EdU signal intensity distribution of individual replicates for RKO IDH2-WT and IDH2-R172K cells. H. Quantification of Mean EdU+ signal intensity from G (3 replicates per condition). I. 2-dimensional plot of EdU signal intensity for RKO IDH2-WT and IDH2-R172K cells with DNA content (Hoechst) in horizontal axis. J. Distribution of EdU signal intensity in EdU+ (solid) and EdU- (hollow) C10T1/2 cells transduced with IDH2-WT and IDH2-R172K expressing vectors. K. 2-dimensional plot of EdU signal intensity for C10T1/2 IDH2-WT and IDH2-R172K cells with DNA content (Hoechst) in horizontal axis. L. Cell cycle distribution frequencies in C10T1/2 IDH2-WT and IDH2-R172K cells. M. 2-dimensional plot of EdU signal intensity for U2OS cells treated with indicated drugs. N. 2-dimensional plot of EdU signal intensity for RKO cells treated with indicated drugs. O. Distribution of EdU signal intensity for RKO cells treated with indicated drugs. P. Cell cycle distribution frequencies for cells in O. Q. 2-dimensional plot of EdU signal intensity and DNA content (Hoechst) for RKO cells treated with indicated drugs. Error bars in B and H represent standard deviation. P values in B and E are indicated and correspond to unpaired t-test.

**Supplementary Figure 4.**
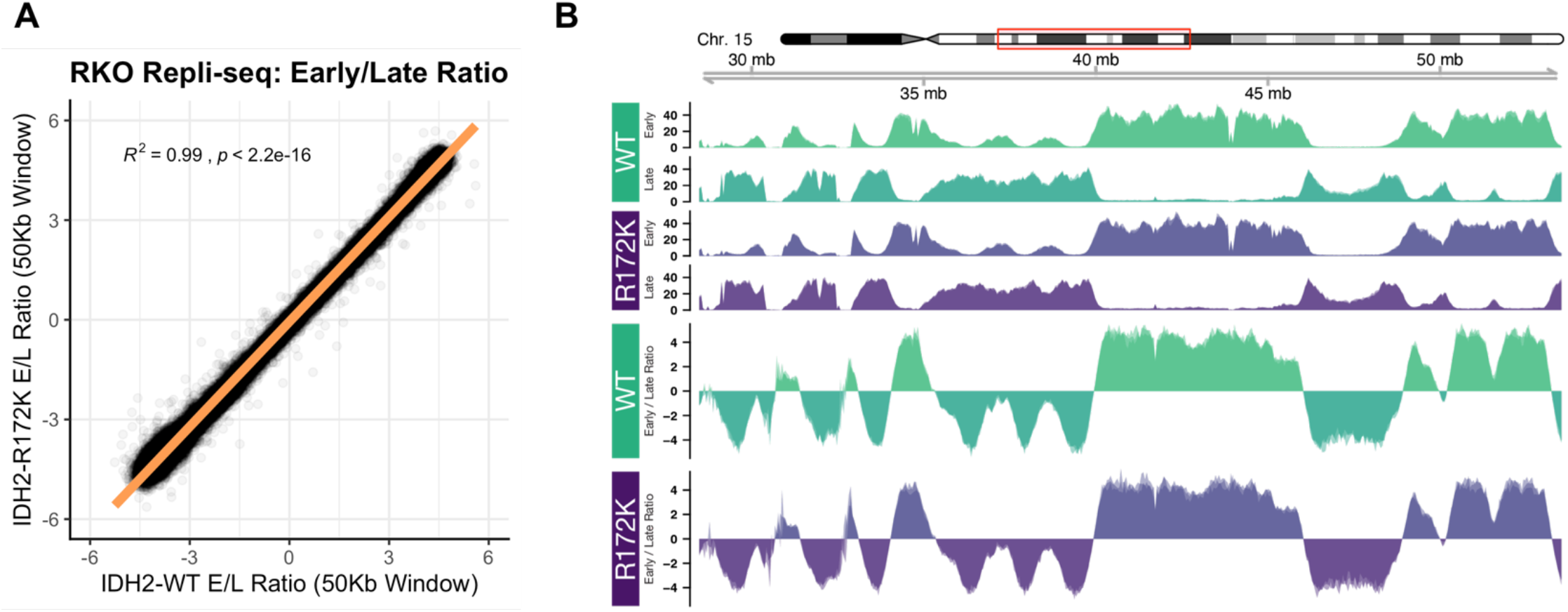
A. Distribution of E/L ratios of reads per 50 Kb windows plotted by IDH status (IDH2-WT, horizontal axis; IDH2-R172K, vertical axis) in RKO cells. B. Early and Late repli-seq tracks from IDH2-WT and IDH2-R172K RKO cells. Ratio of early/late tracks is plotted in lower two sections. In A, R^2^ and p value correspond to the Pearson correlation coefficient.

**Supplementary Figure 5.**
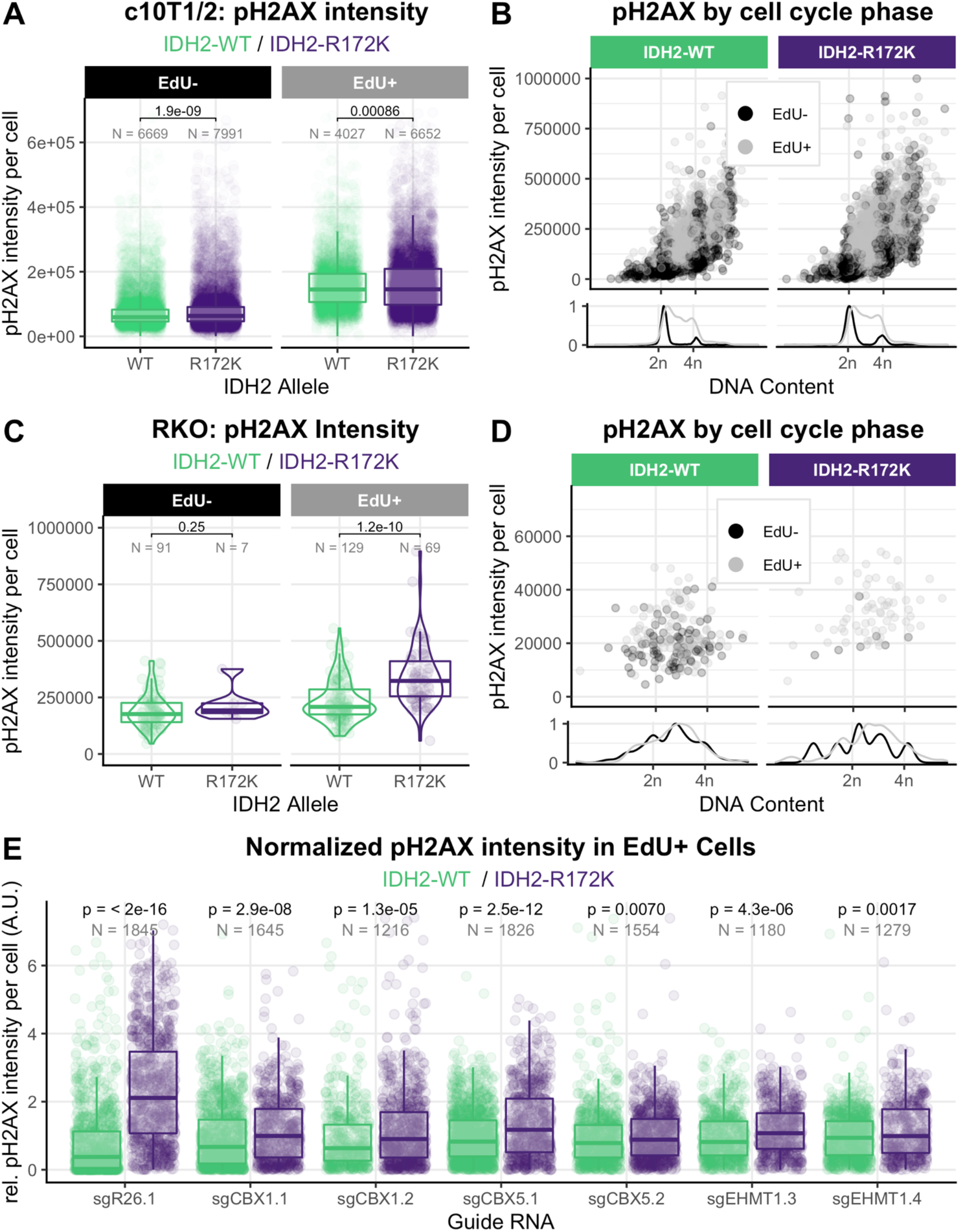
A. Distribution of pH2AX signal intensity in C10T1/2 IDH2-WT and IDH2-R172K cells by EdU signal intensity. B. pH2AX signal intensity from A plotted by EdU signal intensity (EdU-in black, EdU+ in grey) and DNA content determined by quantification of DAPI signal on horizontal axis. Density distribution of DNA content shown in lower panel. C. Distribution of pH2AX signal intensity in RKO IDH2-WT and IDH2-R172K cells by EdU signal intensity. D. pH2AX signal intensity from C plotted by EdU signal intensity (EdU-in black, EdU+ in grey) and DNA content determined by quantification of DAPI signal on horizontal axis. Density distribution of DNA content shown in lower panel. E. Normalized pH2AX signal intensity in U2OS IDH2-WT and IDH2-R172K cells transduced with indicated CRISPR/Cas9 vectors as per Figure 5G, showing additional CRISPR/Cas9 vectors per gene. P values in A, C and E correspond to Wilcoxon tests.

